# Mapping the forest disturbance regimes of Europe

**DOI:** 10.1101/2020.03.30.015875

**Authors:** Cornelius Senf, Rupert Seidl

**Affiliations:** Ecosystem dynamics and forest management group, Technical University of Munich, Hans-Carl-von-Carlowitz-Platz 2, 85354 Freising, Germany; Institute for Silviculture, University of Natural Resources and Life Sciences (BOKU), Vienna, Austria; Berchtesgaden National Park, Doktorberg 6, 83471, Berchtesgaden, Germany

**Keywords:** Forest disturbances, Disturbance regimes, Remote sensing, Forest resilience, Forest management

## Abstract

Forest disturbances shape ecosystem composition and structure, and changes in disturbance regimes can have strong consequences for forest ecosystem services. Yet we currently lack consistent quantitative data on Europe’s forest disturbance regimes and their changes over time. Here we used satellite data to map three decades (1986-2016) of forest disturbances across continental Europe, covering 35 countries and a forest area of 210 million ha at a spatial grain of 30 m, and analyzed the patterns and trends in disturbance size, frequency and severity. Between 1986 and 2016, 17% of Europe’s forest area was disturbed by anthropogenic and/or natural causes. The 25 million individual disturbance patches had a mean patch size of 1.09 ha (range between 1^st^ and 99^th^ percentile 0.18 – 10.10 ha). On average 0.52 (0.02 – 3.01) disturbances occurred per km^2^ every year, removing 77% (22 – 100%) of the canopy. While trends in disturbance size were highly variable, disturbance frequency increased and disturbance severity decreased since 1986. Changes in disturbance rates observed for Europe’s forests are thus primarily driven by changes in disturbance frequency (i.e., more disturbances), and only to a lesser extent by increasing disturbance size. We here present the first continental-scale characterization of Europe’s forest disturbance regimes and their changes over time, providing spatially explicit information that is critical for understanding the ongoing changes in forest ecosystems across Europe.

---

Forests cover 33 % of Europe’s total land area and provide important ecosystem services to society, ranging from carbon sequestration to the filtration of water, protection of soil from erosion and human infrastructure from natural hazards ^1^. Europe’s forests have expanded in recent decades ^2^ and have accumulated substantial amounts of biomass due to intensive post-WWII reforestation programs, changes in management systems, and timber harvesting rates that remained below increment ^3^. This success story of 20^th^ century forestry in Europe, however, also has side effects, as the resultant changes in forest structure and composition have – in combination with climate change – led to an episode of increasing forest disturbances in recent decades ^4–7^. Increasing forest disturbances have the potential to erode Europe’s carbon storage potential ^8,9^ and also impact other important ecosystem services provided by Europe’s forests ^10,11^. Given a predicted increase in the demand for wood ^1^ and an expected future intensification of forest dieback under climate change ^12^, it is fundamental to both understand and increase the resilience of Europe’s forests to changing disturbances ^13–15^.

Understanding the ongoing changes in forest ecosystems and developing management strategies to increase their resilience requires a robust quantitative understanding of the prevailing disturbance regimes ^16,17^. Disturbance regimes characterize the cumulative effects of all disturbance events occurring in a given area and time period, and are often characterized by metrics such as the size, frequency, and severity of disturbances occurring in a given area ^16^. In Europe, forests have been utilized by humans for centuries, transforming species composition and structure ^18–20^, and consequently also the natural disturbance regimes of forests. In addition to this indirect effect, human land-use is directly disturbing forest canopies through timber harvesting, altering the rate and spatial patterns of forest disturbances compared to natural systems ^21^. Human land-use also interacts with natural disturbances, e.g. by salvage logging of disturbed timber ^22^ and shortening early seral stages through planting ^23^. More broadly, forest management alters biological legacies and landscape structure ^22,24^, with feedbacks on subsequent disturbances. Due to the intricate linkages between natural and human processes driving forest disturbances in Europe, characterizing the disturbance regimes of Europe’s forests requires a holistic perspective covering both natural and human disturbances.

For Europe, there is little quantitative information on disturbance regimes and their changes through time available to date, especially if considering both natural and human disturbances. While previous studies have characterized the disturbance regimes of some of Europe’s forest ecosystems ^4,18,25–27^, those studies have either focused on purely natural processes, lack a spatially and temporally consistent data source, or focus only on the regional scale. Due to this lack of quantitative information at continental scale, we do, for instance, not know how disturbance size, frequency and severity vary across Europe. Furthermore, while recent studies indicate an increase in disturbance rates across Europe’s natural and managed forests ^4,6^, it remains unknown whether this change is mainly the result of changes in disturbance frequency (i.e., more disturbance events) or disturbance size (i.e., larger individual disturbance patches). Likewise, our quantitative knowledge of changes in disturbance severity is scant, and it remains unclear whether disturbances in Europe have become more severe in recent decades (e.g., through increased burn severities ^28^) or whether recent changes in forest management approaches (e.g., the adoption of “close-to-nature” silviculture ^29^) have reduced disturbance severity, as reported for parts of Central Europe ^4^, for instance.

Here, our aim was to map and characterize the disturbance regimes of Europe’s forests 1986 – 2016. Our specific research questions were: (I) What is the size, frequency and severity of forest disturbances across Europe’s forests? (II) How did size, frequency and severity of forest disturbances change over the past three decades? We addressed these two questions by mapping forest disturbance occurrence and severity continuously for continental Europe (35 countries covering 210 million ha of forest) at a spatial grain of 30 m using more than 30,000 satellite images and nearly 20,000 manually interpreted reference plots. Subsequently, we characterized both the spatial variation of disturbance size, frequency and severity and their temporal trends over time at the continental scale, thus providing the quantitative baseline critically needed for understanding current changes in Europe’s forest ecosystems.

## Results

### Disturbance maps

We identified a total of 36 million individual disturbance patches occurring across Europe between 1986-2016, equaling a disturbed forest area of 39 million ha or 17 % of Europe’s forest area (Figure 1). The overall accuracy of the map was 92.5 ± 2.1 % (mean ± standard error), with a commission error of 14.6 ± 1.8 % and an omission error of 32.8 ± 0.3 % for detecting disturbances (see Supplementary Table S2). Omission errors were mainly related to low severity disturbances that could not be separated from noise (Figure S4). The mean absolute error between the estimated disturbance year and the manually interpreted disturbance year was three years (Figure S5), with 77 % of the assigned disturbance years being within three years of the manually interpreted disturbance year. We derived a continuous value ranging from zero to one as measure of disturbance severity (see Figure 1C). The severity measure expresses the probability of a disturbance being stand-replacing, with zero indicating no change in the dominant canopy and one indicating a complete removal of the forest canopy in a disturbance. The disturbance severity measure was well able to differentiate between un-disturbed areas (no loss of forest canopy), non-stand-replacing disturbances (partial loss of forest canopy), and stand-replacing disturbances (complete loss of forest canopy; Figure S6), and thus well represents the variable disturbance severities prevailing across Europe’s forests.

**Fig. 1:**
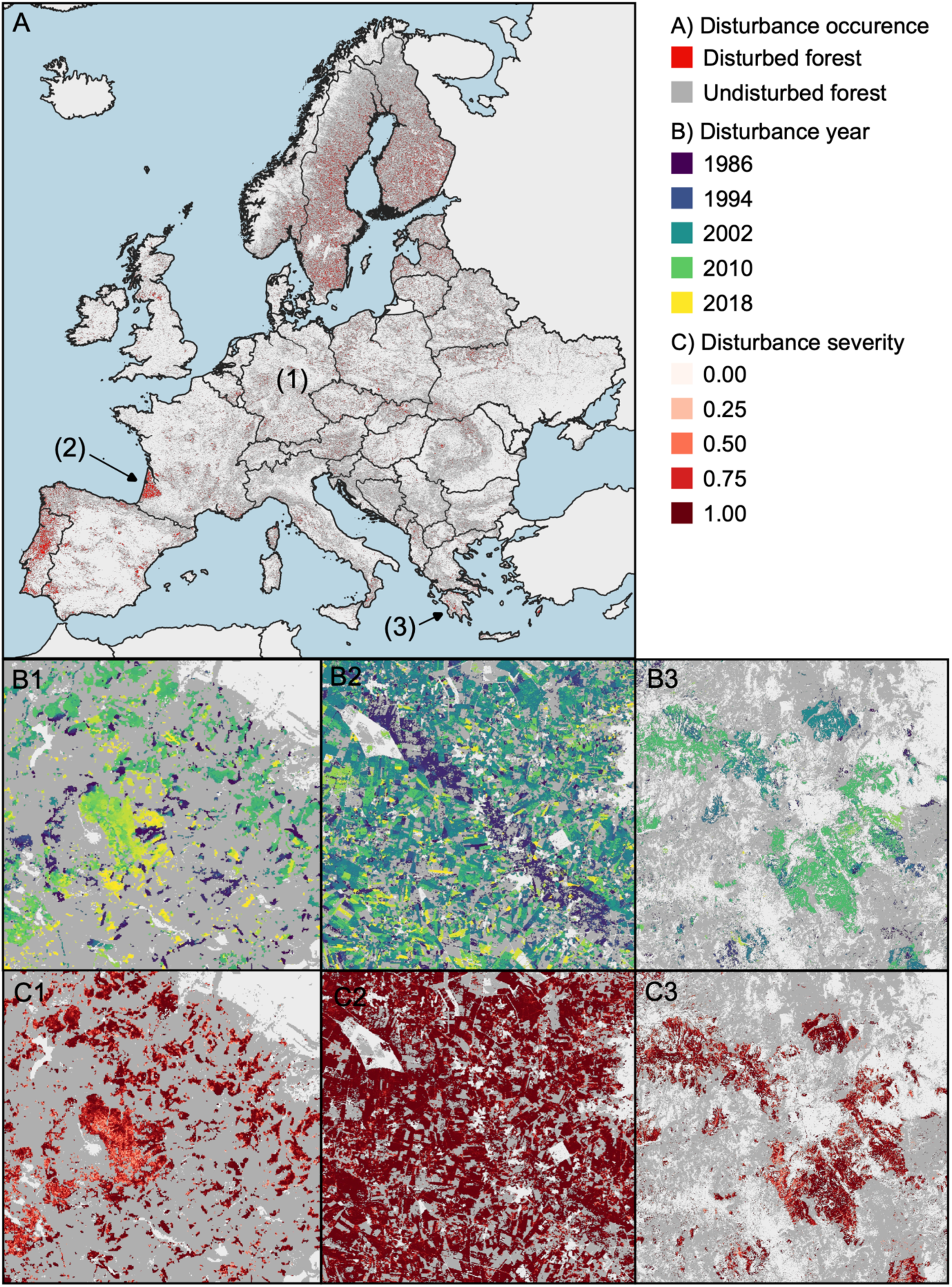
Forest disturbances in Europe 1986-2016. Disturbance maps were derived from analyzing more than 30,000 Landsat images across continental Europe. Panel A shows the occurrence of disturbances across Europe. Panels B show the disturbance year and panels C show the disturbance severity for three selected areas: (1) A bark beetle outbreak of varying severity in and around Harz National Park (Germany); (2) salvage-logged wind disturbance in an intensively managed plantation forest in the Landes of Gascony (France) with very high disturbance severity; and (3) fire disturbances on the Peloponnese peninsula (Greece), with variable burn severity. See Supplementary Figure S10 for a high-quality version of the main disturbance map.

### Disturbance regimes

The average patch size of forest disturbances was 1.09 ha, but the disturbance size distribution was highly left-skewed (Figure 2-B). The median disturbance size was only 0.45 ha, with 78 % of the disturbances being smaller than 1 ha and 99 % of the disturbances being smaller than 10 ha (Table 1). The largest disturbance patch mapped across Europe was a 16,617 ha large forest fire occurring in 2012 in southern Spain. The average disturbance frequency was 0.52 patches per km^2^ of forest area per year (median of 0.37 patches per km^2^), with highest frequencies (highest 1 %) ranging from 3 to 31 patches per km^2^ (Table 1). Disturbance severity ranged from 0.23 to 1.00, with an average of 0.77 (median of 0.83). That is, more than half of the disturbed patches across Europe had a very high probability of being stand replacing, indicating a high prevalence of high severity disturbances.

**Table 1:**
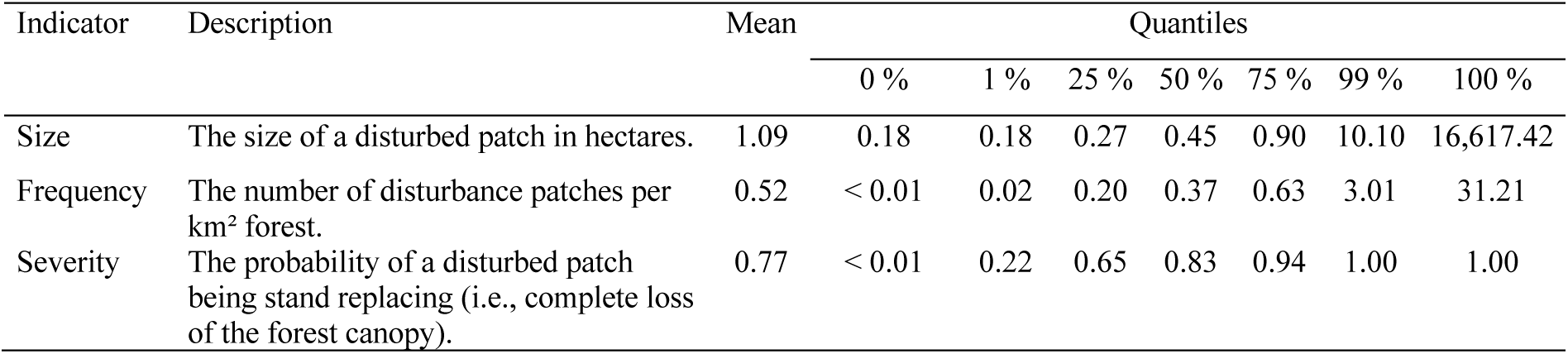
Distribution of the size, frequency and severity of disturbances across Europe’s forests (see Supplementary Table S3 for values by country).

**Fig. 2:**
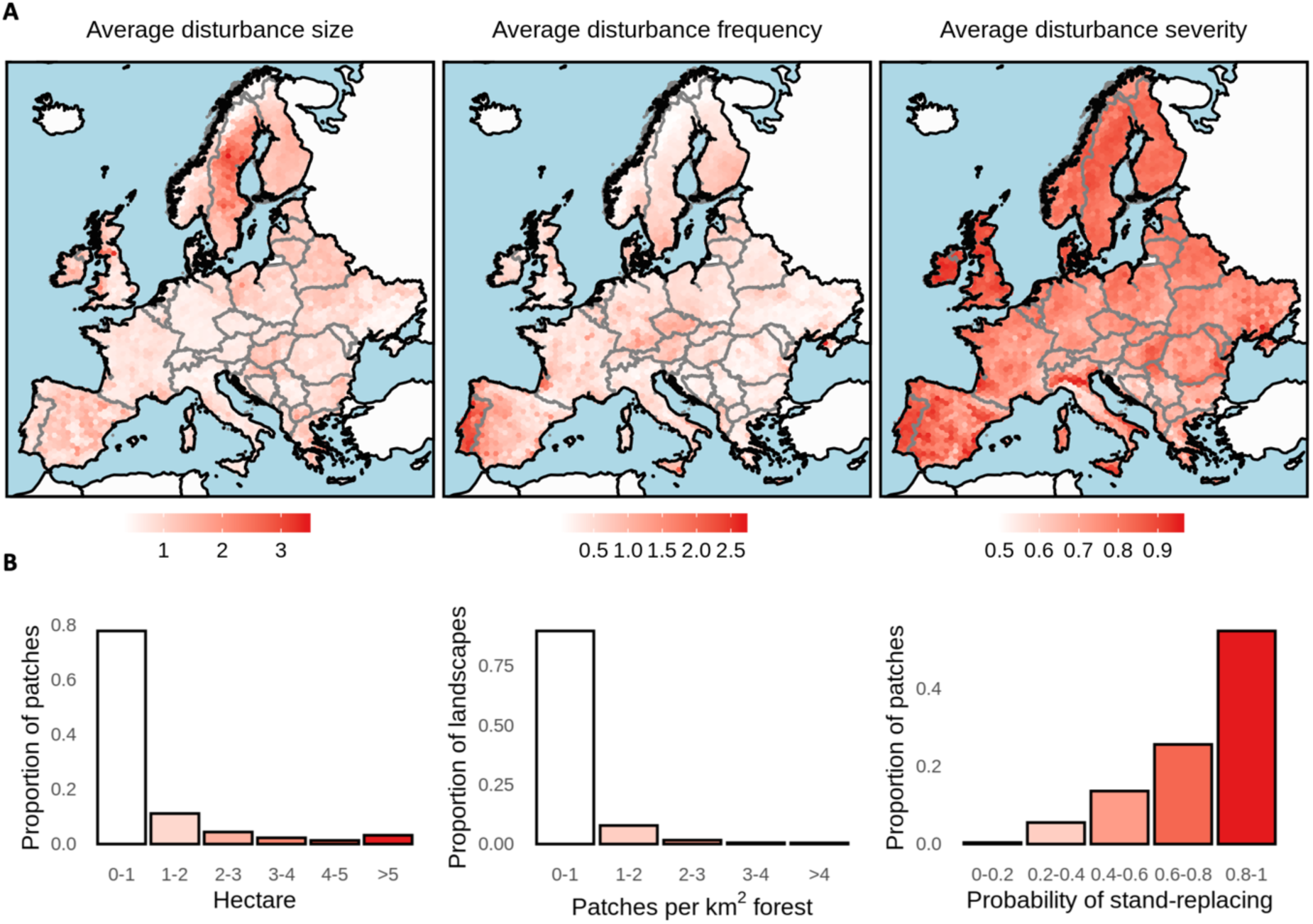
(A) Maps of average disturbance size, frequency and severity calculated for hexagons on a 50 km grid across continental Europe. (B) Distribution of average disturbance size, frequency and severity across Europe.

Spatial variability in the size, frequency and severity of forest disturbances is high across Europe (Figure 2). Disturbance patches are generally larger in Northern and Southern Europe compared to Central Europe. Also, Eastern Europe has larger disturbance patches compared to Western Europe (Figure 2). Above-average disturbance frequencies were found in Central Europe, the hemi-boreal zone, parts of France and the Iberian Peninsula (Figure 2). The highest disturbance frequencies (i.e., > 3 patches per km^2^) occurred almost exclusively in Portugal. Disturbance severity was more evenly distributed than the other two disturbance regime indicators (Figure 2), with a tendency towards higher severities in the Atlantic forests of Ireland and the United Kingdom, the Iberian Peninsula, the Po-Valley in Italy, and the Pannonian Basin. In contrast, low disturbance severities were recorded for South-Eastern Europe along the Dinaric mountain range, as well as in the Apennine mountains of Italy.

### Trends in disturbance regimes

Disturbance regimes changed profoundly between 1986 and 2016, but trends differed with disturbance regime indicator (Figure 3). Changes in disturbance size were variable across Europe. Hot spots of increasing disturbance size were in the Baltic states, the United Kingdom, Ireland, and Italy (Figure 3), whereas trends were largely negative in Eastern Germany, western Poland and southeastern Europe (Figure 3). Disturbance frequency showed a more consistent increase than disturbance size, with disturbance frequency increasing on 74 % of Europe’s forest area (Table 2). Hot spots of increasing disturbance frequency were located in Central and Eastern Europe (Figure 3), whereas negative trends in disturbance frequency were found for Belarus, Albania and Greece as well as parts of western Europe and northern Fennoscandia (Figure 3). In contrast, disturbance severity decreased for 88 % of the European forest area (Table 2), with particularly strong negative trends in Central and Southeastern Europe (Figure 3).

**Table 2:** Distribution of the trends in size, frequency and severity of disturbances across Europe’s forests 1986 – 2016 (see Supplementary Table S3 for values by country).

| Indicator | Mean (across disturbed patches) | Mean (weighted by forest area) | Proportion of forest area with positive trends | Proportion of forest area with no trend |
| --- | --- | --- | --- | --- |
| Size (ha) |  |  |  |  |
| Mean | 0.41 | 0.33 | 0.65 | 0.00 |
| 50 % quantile | 0.21 | 0.23 | 0.19 | 0.78 |
| 75 % quantile | 0.53 | 0.53 | 0.54 | 0.32 |
| 100 % quantile | 0.35 | 0.15 | 0.52 | 0.00 |
| Frequency (# per km<sup>2</sup> forest area) | 1.17 | 1.19 | 0.74 | 0.01 |
| Severity (0-1) |  |  |  |  |
| Mean | -0.31 | -0.33 | 0.12 | 0.00 |

**Fig. 3:**
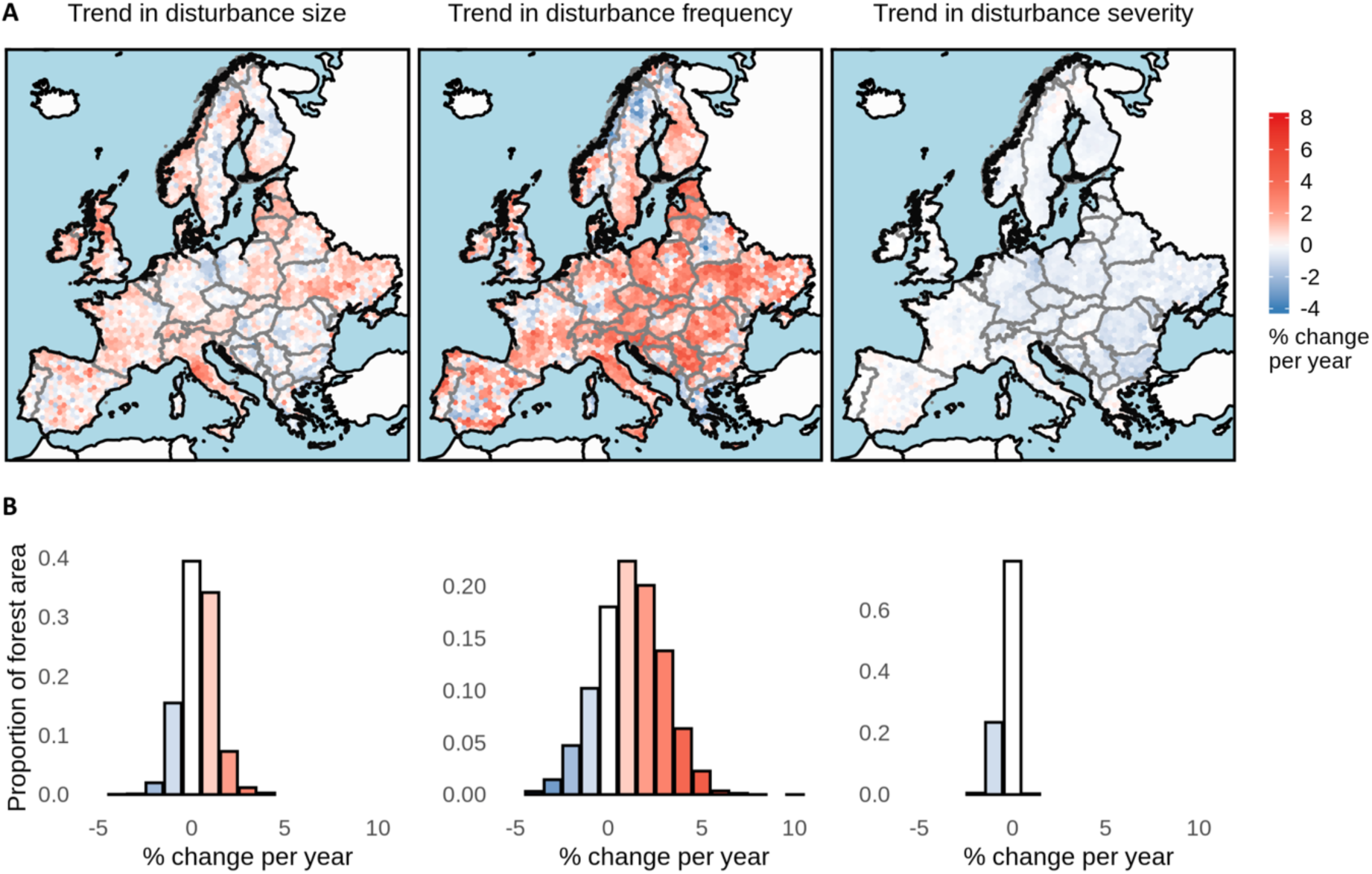
Trends in Europe’s forest disturbance regimes 1986 – 2016. (A) Maps of trends in disturbance size, frequency and severity calculated at a 50 km hexagon grid across continental Europe. (B) Distribution of forest area among trend classes.

While the mean disturbance size generally increased across Europe (65 % of Europe’s forests had an increasing trend in mean disturbance size; Table 2) the median disturbance size was more stable (increasing for only 19 % of Europe’s forests; Table 2). Hence, the disturbance size distribution widened over time, with an increase in large disturbance patches (i.e., in the 75 % quantile and maximum of the disturbance patch size distribution; Table 2) in approximately half of Europe’s forests. Overall, changes in disturbance frequency explained 71 % of the variability in changing disturbance rates (i.e., the trend in the annual percent of forest area disturbed), whereas changes in disturbance size only accounted for 24 % (Supplementary Figure S7). Thus, changes in forest disturbance rates observed in Europe are primarily driven by more frequent disturbances, and only to a lesser extent by increasing disturbance sizes.

## Discussion

We here provide the first quantitative and spatially explicit characterization of Europe’s forest disturbance regimes, highlighting the wide variety in disturbance sizes, frequencies and severities prevailing across the European continent. While forest type and general biophysical environment certainly explain part of the variability in the disturbance regimes of Europe ^6,30^, it is likely also the variability in forest management approaches across Europe’s forests that plays a fundamental role in explaining the observed patterns. Forest management approaches across Europe range from small-scale approaches aiming for continuous forest cover to even-aged forestry based on clearfelling and high-intensity short rotation systems ^31,32^. For example, many countries that predominantly use small-scale management approaches (e.g., Slovenia, Switzerland ^33^) were characterized by substantially smaller disturbances sizes and lower disturbances severities in our data (Supplementary Figure S8), despite also experiencing large-scale natural disturbances ^34^. This clearly contrasts with countries largely apply even-aged forest management approaches (e.g., Finland, Sweden ^35^) or have high shares of plantation forests (e.g., Denmark, Hungary, Ireland ^1^), which have on average larger patch sizes and higher disturbance severities (Supplementary Figure S8). Disturbance regimes thus varied widely between countries, reflecting differences in management objectives and management histories ^36^. In some instances, these differences occur even for countries which have very similar forest types and biophysical environments, which would suggest a comparable natural disturbance regime (see, e.g., Figure S9 for additional examples). A substantial part of the spatial variability in disturbance size, frequency and severity observed here are thus likely driven by variation in forest management across the European continent.

The disturbance regimes of Europe’s forests are changing profoundly. We here show that the previously reported increase in disturbance rates ^4,7^ is primarily an effect of increasing disturbance frequency, while disturbance patch size distributions are becoming more variable and disturbance severities are decreasing. The strong increase in disturbance frequencies might be caused by both increasing wood production and increasing natural disturbances reported throughout Europe ^9^, with both factors likely interacting (i.e., increasing natural disturbances triggering increased salvage harvesting). The widening of the patch size distribution likely results from the combined effects of changes in management approaches towards smaller intervention sizes (i.e., single-tree or group selection ^29^) and simultaneously increasing natural disturbance activity, leading to infrequent but large canopy removals (e.g., large-scale storm events ^37^ or large-scale fires ^38^). The combined effects of changes in management approaches and natural disturbances also likely explains continental-scale decreases in disturbance severity, as management systems are increasingly optimized to reduce impact ^39^ and many natural disturbances that occur frequently (i.e., bark beetle infestations and small-scale windthrow) are characterized by mixed severities ^40^.

We here provide the first high-resolution forest disturbance map for continental Europe covering three decades of forest development, a dataset of importance for future research on the dynamics of Europe’s forests. Yet, there are methodological limitations that should be considered when using the data presented here. First, we do not distinguish disturbance agents in our analysis, that is an individual disturbance patch can currently not be attributed to either natural or human origin. While methodological advances in attributing disturbance agents based on satellite-based forest change products have been made recently ^41^, those approaches are not yet applicable at the spatial and temporal scale of our analysis. The key reasons for this are missing reference data on the actual occurrence of disturbance agents and the fact that management signals are often superimposed on natural disturbances (i.e., subsequent salvage logging). Future work should thus aim for improved attribution algorithms that consider the complex interactions between humans and natural processes in Europe’s forest ecosystems more explicitly. Second, we here only map the greatest disturbance per pixel, that is there is only one disturbance event recorded for the whole 30-year period for each 30 × 30 m pixel. For short-rotation systems we thus might miss some disturbances if, e.g., two harvests have occurred in the past three decades. Finally, we note that despite careful processing (historic) satellite data can be noisy, preventing the detection of very low-severity disturbances. This limitation is intrinsic to the data used herein, there are, however, very few alternative data sources that allow the consistent analysis of vegetation dynamics over three decades at continental scale. Despite these limitations we are confident that our first quantitative and spatially explicit analysis of patterns and trends in forest disturbances provide a crucial step towards better understanding the ongoing changes in Europe’s forest ecosystems.

## Materials and Methods

### Reference data

Acquiring consistent reference data across large areas – such as continental Europe – is challenging. We here made use of manually interpreted satellite data, serving as valuable alternative to field-based data ^42^. Manual interpretation of satellite data for calibrating and validating Landsat-based forest change maps is a well-established approach and has been used in numerous studies previously ^43–46^. In essence, an interpreter inspects the temporal profile of the spectral trajectory of a Landsat pixel and, with the help of Landsat image chips and very high-resolution imagery available in Google Earth, makes a well-informed call whether the trajectory represents stable forest canopy cover or whether a mortality event occurred ^47^. We here used a previously established set of 19,996 interpreted Landsat pixels ^7^ as reference data. The initial sample was drawn at random within forests of Europe, with samples stratified by country (500 samples per country). As interpreters might declare a plot as no-forest during interpretation (caused by errors in the automatically generated forest mask used as the basis for stratified sampling), the realized sample size varied between countries (Table S1). The response design followed well-documented protocols developed and published previously ^4,7^. Manual interpretation was done by a total of nine interpreters using established software tools ^47^, and the data is freely accessible under following repository: https://doi.org/10.5281/zenodo.3561925

The reference sample set only consisted of forest pixels and there was thus need for substituting the sample with non-forest reference pixels. We therefore drew a country-stratified sample of non-forest pixels using a Landsat-based land cover map from ^48^. Each countries sample size was chosen to match the forest proportion of the respective country (based on data from the FAOSTATS database), that is the total sample of each country equaled a random sample across its terrestrial forested and non-forested land surface (see Table S1). In total we drew 46,461 non-forest reference pixel that, paired with the 19,996 forest reference pixels manually interpreted, totaled to 66,457 reference pixels used for calibration and validation. From the full reference sample, we randomly drew a sub-sample of 5,000 pixels for map validation, and the remaining 61,457 pixels were used for model calibration. The validation sub-sample was drawn proportionally to the size of each country to ensure a consistent and unbiased estimation of mapping accuracies for the final European map product.

### Mapping disturbances

At the core of our mapping workflow we rely on an established time-series segmentation approach called LandTrendr ^49^, implemented in the high-performance cloud-computing environment Google Earth Engine ^50^. In essence, LandTrendr segments annual Landsat pixel time series into linear features, for which a set of metrics can be extracted. We here do not provide details on the underlying LandTrendr routines but focus on the salient details of our mapping workflow (cf. Figure S2 for a graphical outline). The workflow was based on code published in Kennedy et al. ^50^.

In a first step we screened all available Tier 1 Landsat 4, 5, 7 and 8 images in the United States Geological Survey archive. Tier 1 images are delivered as ready-to-use surface reflectance images including a cloud mask, yet we used coefficients from Roy et al. ^51^ to spectrally align the varying sensor types used onboard Landsat 4/5 (Thematic Mapper), Landsat 7 (Enhanced Thematic Mapper Plus), and Landsat 8 (Operational Land Imager). After spectral alignment we filtered all available images for summer-season acquisition dates (1^st^ of June to 30^th^ September) and built annual medoid composites following Flood ^52^.

Second, we ran LandTrendr for two spectral bands (shortwave infrared I and II) and two spectral indices commonly used for forest disturbance and mortality mapping ^45,46,53–55^: the Tasseled Cap wetness (TCW) and the Normalized Burn Ration (NBR). We used a standard parameter set for LandTrendr with no filtering or thresholding and thus allowing for maximum sensitivity in detecting change (i.e., allowing for a high commission error).

Third, we extracted the greatest change segment from each pixel’s LandTrendr trajectory, fit to both spectral bands and both spectral indices. From the greatest change segment we derived a set of three metrics describing the magnitude, duration and rate of change ^53^ as well as a measure of the signal-to-noise ratio as described in Cohen et al. ^54^. We further derived the spectral band/index value prior to, and the rate of change following the greatest change segment. Similar metrics as the ones used here have been applied also in previous studies mapping forest cover changes ^44,46,55^.

Fourth, we used the set of metrics derived from the greatest change segment for the two spectral bands and the two spectral indices, the calibration data outlined in the previous section, and random forest classification ^56^ to classify each pixel into either no-forest, undisturbed forest, or disturbed forest (i.e., at least one disturbance event during the study period). This last step filtered out commission errors by LandTrendr and thus greatly improves mapping accuracy compared to purely automatic algorithms ^57^. Yet, we experienced difficulties in correctly separating forest and no-forest areas solely based on LandTrendr outputs. This was due to high spectral changes in agricultural areas, which were identified as disturbances by LandTrendr. To tackle this problem, we added a three-year Tasseled Cap Brightness, Greenness and Wetness median composite centered on 1985 and 2018, respectively, to the classification stack. The additional six bands delivered more detailed spectral information on stable forest and non-forest pixels. Finally, we applied the trained random forest model to the full classification stack (i.e., LandTrendr metrics from the two spectral bands and two spectral indices plus the Tasseled Cap composite from 1985 and 2018) to consistently map the categories no forest, undisturbed forest and disturbed forest across continental Europe. We validated the final map using the validation sub-sample described in the previous section. We derived a confusion matrix and report overall accuracy, errors of commission and errors of omission following best-practice recommendations given in ^42^.

Fifth, while the thus derived map indicates whether a mortality event has happened or not, it does not inform about when the mortality event happened. We therefore calculated the year of the disturbance onset (i.e., the year of the greatest spectral change) from all spectral bands and spectral indices using an automated majority vote. If there was a tie (e.g., all four bands/indices indicated a different year), we reverted to the median value. To validate this processing step, we compared the year assigned from LandTrendr to the manually interpreted year of disturbance for the 19,996 forest reference plots.

### Spatial filtering

The last step in creating disturbance maps for continental Europe was to apply a set of spatial filters for removing unrealistic outliers from the resulting disturbance maps and enhancing spatial pattern analysis. We first set a minimum mapping unit of two 30 × 30 m pixels (i.e., 0.18 ha) and removed all disturbance patches smaller than the minimum mapping unit. In a second filtering step, we identified all patches smaller than the minimum mapping unit for each year, and assigned them to the year of the surrounding disturbed pixels (if any), thus accounting for artefacts related to uncertainties in the correct identification of the disturbance year (see Figure S3). In a final filtering step, we removed holes within disturbance patches smaller than the minimum mapping unit by filling them with the year of the surrounding pixels. While the filtering was done to improve the spatial analyses described in the following section, we note that the filtering was applied after the accuracy assessment. The accuracy assessment thus reports the raw classification performance without additional filtering.

### Characterizing disturbance regimes and their changes

From the annual forest disturbance maps we calculated three disturbance regime indicators based on Turner ^16^ and Johnstone et al. ^17^: disturbance size, frequency and severity. Disturbance size and severity were calculated at the patch level and subsequently aggregated to the landscape level, while disturbance frequency was calculated at the landscape level directly. Disturbance size is the number of disturbed pixels for each individual patch (patches were defined annually using rook-contiguity) multiplied by pixel size (0.09 ha). For calculating disturbance frequency, we sub-divided the total study area into a 50 * 50 km hexagon grid (here representing the landscape scale, hexagon area of 2165 km^2^), with a total number of 3,240 hexagons across Europe’s land area. We chose hexagons over squares as hexagons minimize the spatial differences to the more complex landforms of the European continent and the borders of European countries ^58^. For each hexagon, we counted the number of individual disturbance patches per year and divided this number by the total forest area within the hexagon, resulting in a measure of the number of disturbed patches per km^2^ forest area per year as an indicator of disturbance frequency.

For quantifying disturbance severity, we made use of the spectral change magnitude provided by the LandTrendr analysis. The spectral change magnitude is well correlated with changes in forest structure during disturbance ^45,53,59^ and we here use it as proxy of disturbance severity. To combine and scale the spectral change magnitude from all four spectral bands/indices into one measure of disturbance severity we used logistic regression to predict the occurrence of stand-replacing disturbances from the four spectral change magnitudes. Data on stand replacing disturbances were generated from the reference sample by analyzing the manually interpreted land cover after a disturbance segment. If the land cover switched to non-treed following a disturbance segment (e.g., after clear-cut harvest or high intensity fire), the disturbance was assumed to be stand-replacing. If the land cover remains treed following a disturbance segment (e.g., following a thinning operation or a low intensity windthrow), the disturbance was classified as non-stand-replacing. The method is based on Senf et al. ^4^ who showed that visual interpretation of post-disturbance land cover is an accurate measure for separating stand-replacing from non-stand-replacing disturbances. By predicting the occurrence of stand-replacing disturbances (i.e., complete removal of the canopy and thus a disturbance with very high severity), we scale the spectral change magnitudes to a value between zero and one, where one indicates complete loss of the canopy (i.e., high severity disturbance) and values close to zero indicate little change in the forest canopy (i.e., low severity disturbance). Values in-between represent variable levels of canopy loss and thus intermediate disturbance severities. While it is difficult to validate this proxy retrospectively across Europe (i.e., no reliable pan-European data on canopy changes during past disturbance events exists), we here performed an indirect validation by comparing the disturbance severity measure among stand-replacing disturbances, non-stand-replacing disturbances and undisturbed reference pixels (see Supplementary Figure S6).

For spatially visualizing disturbance size, frequency and severity as well as for calculating and visualizing trends we aggregated patch-based metrics (i.e., disturbance size and severity) to the landscape level (i.e., the hexagon) by calculating the arithmetic mean. We report the mean over the median as it is sensitive to changes in both the central tendency and the spread of the distribution, but we also include other descriptors in the Tables and Supplement. Trends in disturbance size, frequency and severity were quantified using a non-parametric Theil–Sen estimator, which is a non-parametric measure of monotonic trends in time series insensitive to outliers ^60^.

## Acknowledgements

C. Senf acknowledges funding from the Austrian Science Fund (FWF) Lise-Meitner Program (Nr. M2652). R. Seidl acknowledges funding from FWF START grant Y895-B25. We thank Justin Brasten (Oregon State University) for making the code of LandTrendr open source, which greatly helped in implementing this research. We furthermore are grateful to three anonymous reviewers for constructive comments on an earlier version of the manuscript.

## Author contribution

CS and RS designed the research; CS performed all computations and analyses; CS wrote the manuscript with input from RS.

## Data availability

All maps presented in this research paper will be made publicly available after peer-review.

## Code availability

The code used for processing the Landsat data is available at https://github.com/eMapR/LT-GEE. The research code used for creating the maps will be made available with the final published version of this manuscript.

