## Supplementary Materials for "Mapping the forest disturbance regimes of Europe"

Cornelius Senf and Rupert Seidl

**Table S1:** Number of reference plots and their distribution across countries.

| Country | Forest | No-Forest | Total | Validation | Calibration |
|---|---|---|---|---|---|
| Albania | 415 | 1132 | 1547 | 25 | 1522 |
| Austria | 1828 | 2050 | 3878 | 73 | 3805 |
| Belarus | 442 | 700 | 1142 | 181 | 961 |
| Belgium | 334 | 1161 | 1495 | 27 | 1468 |
| Bosnia and Herzegovina | 393 | 391 | 784 | 44 | 740 |
| Bulgaria | 418 | 863 | 1281 | 96 | 1185 |
| Croatia | 433 | 558 | 991 | 50 | 941 |
| Czech Republic | 1401 | 2847 | 4248 | 69 | 4179 |
| Denmark | 401 | 2452 | 2853 | 38 | 2815 |
| Estonia | 377 | 365 | 742 | 39 | 703 |
| Finland | 450 | 202 | 652 | 294 | 358 |
| France | 413 | 507 | 920 | 477 | 443 |
| Germany | 1227 | 2615 | 3842 | 311 | 3531 |
| Greece | 412 | 956 | 1,368 | 109 | 1,259 |
| Hungary | 349 | 1,199 | 1,548 | 81 | 1,467 |
| Ireland | 298 | 2,474 | 2,772 | 61 | 2,711 |
| Italy | 363 | 660 | 1,023 | 262 | 761 |
| Latvia | 389 | 483 | 872 | 56 | 816 |
| Lithuania | 416 | 858 | 1,274 | 56 | 1,218 |
| Moldova | 302 | 2,803 | 3,105 | 29 | 3,076 |
| Montenegro | 403 | 485 | 888 | 12 | 876 |
| Netherlands | 322 | 2,782 | 3,104 | 31 | 3,073 |
| North Macedonia | 439 | 647 | 1,086 | 22 | 1,064 |
| Norway | 394 | 621 | 1,015 | 271 | 744 |
| Poland | 1,312 | 3,234 | 4,546 | 271 | 4,275 |
| Portugal | 387 | 691 | 1,078 | 77 | 1,001 |
| Romania | 436 | 1,058 | 1,494 | 207 | 1,287 |
| Serbia | 334 | 750 | 1,084 | 77 | 1,007 |
| Slovakia | 1,820 | 2,640 | 4,460 | 43 | 4,417 |
| Slovenia | 412 | 251 | 663 | 18 | 645 |
| Spain | 326 | 557 | 883 | 434 | 449 |
| Sweden | 439 | 265 | 704 | 391 | 313 |
| Switzerland | 1,128 | 2,581 | 3,709 | 36 | 3,673 |
| Ukraine | 451 | 2,115 | 2,566 | 519 | 2,047 |
| United Kingdom | 332 | 2,508 | 2,840 | 213 | 2,627 |
| **Sum** | **19,996** | **46,461** | **66,457** | **5,000** | **61,457** |

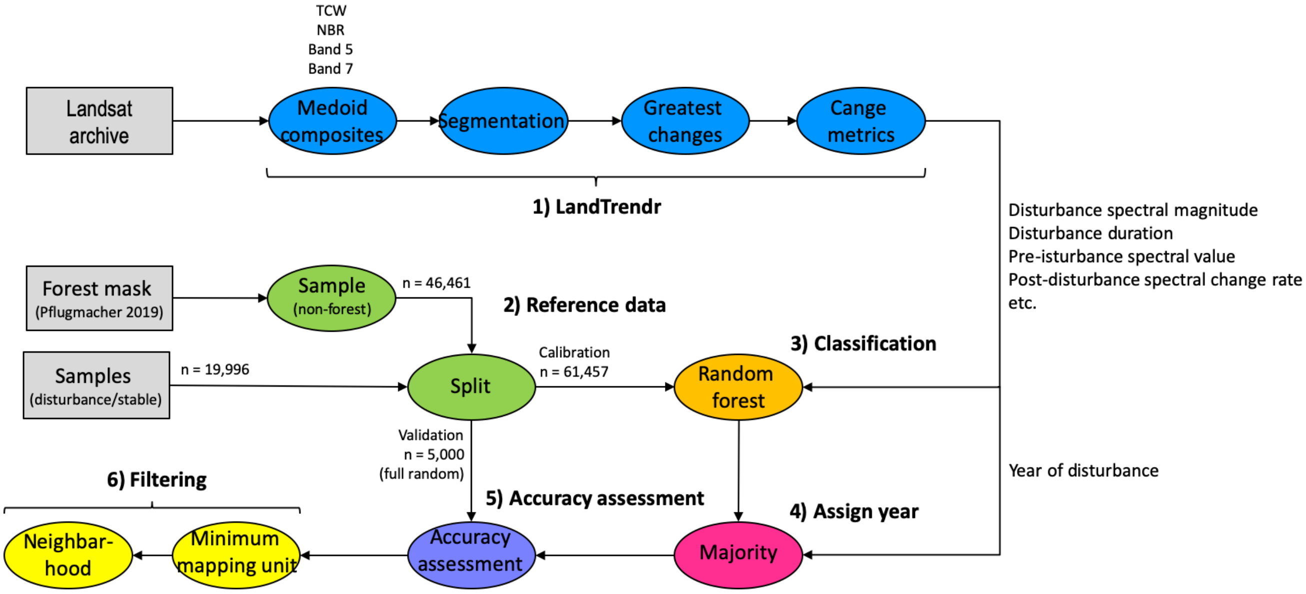


**Figure S2:** Workflow for mapping no-forest areas, undisturbed forests and disturbed forests across continental Europe.

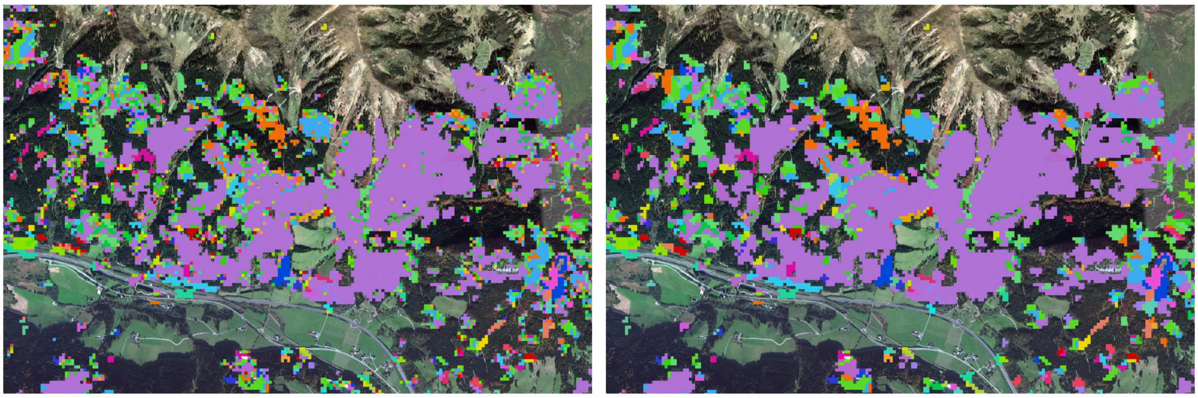

**Figure S3:** Example of the spatial filter applied to the disturbance maps. Colors indicate different years of disturbance.

**Table S2:** Confusion matrix (expressed as proportions), overall accuracy (± standard error) and errors of omission and commission (± standard error), all derived from and independent and randomly distributed validation sample of n = 5,000.

| Mapped | Interpreted | | | Total | Commission error rate |
|---|---|---|---|---|---|
| | Disturbance | Forest | Non-forest | | |
| Disturbance | 0.063 | 0.010 | 0.001 | 0.074 | 0.146 ± 0.018 |
| Forest | 0.026 | 0.236 | 0.007 | 0.269 | 0.121 ± 0.009 |
| Non-forest | 0.005 | 0.026 | 0.626 | 0.657 | 0.048 ± 0.004 |
| Total | 0.094 | 0.272 | 0.634 | 1.000 | |
| Omission error rate | 0.328 ± 0.003 | 0.132 ± 0.004 | 0.012 ± 0.003 | | Overall error rate: 0.075 ± 0.021 |

For **Table S3** see Excel file attached to the manuscript.

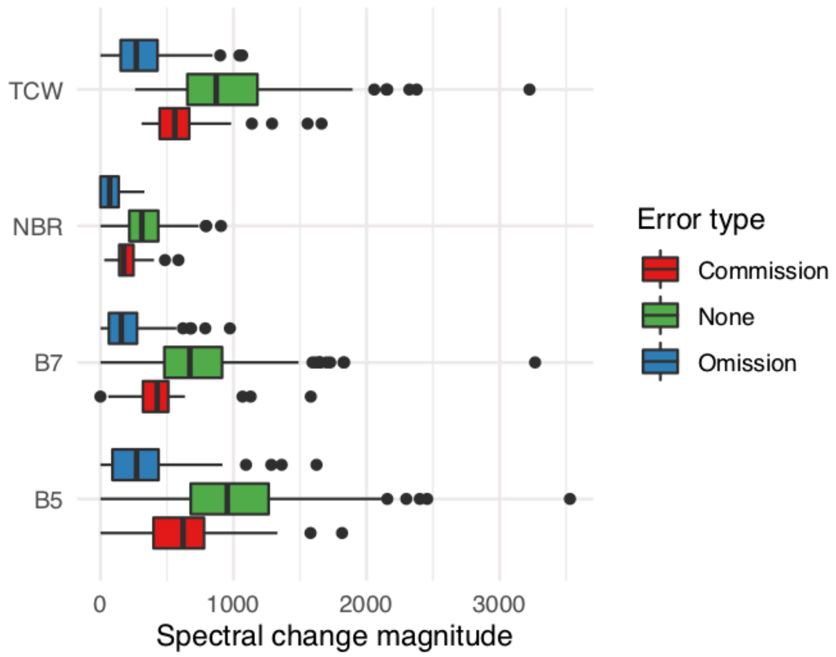


**Figure S4:** Spectral change magnitude in Tasseled Cap Wetness (TCW), Normalized Burn Ration (NBR), Landsat shortwave-infrared I (B5), and Landsat shortwave-infrared II (B7) for all reference pixels (n = 5,000) with commission errors, omission errors and no error (i.e., matching label between mapped and interpreted). For omission errors, spectral change magnitudes were substantially lower than for disturbances, highlighting that many omission errors stem from very low spectral changes, indistinguishable from noise in currently available Landsat-based time series methods.

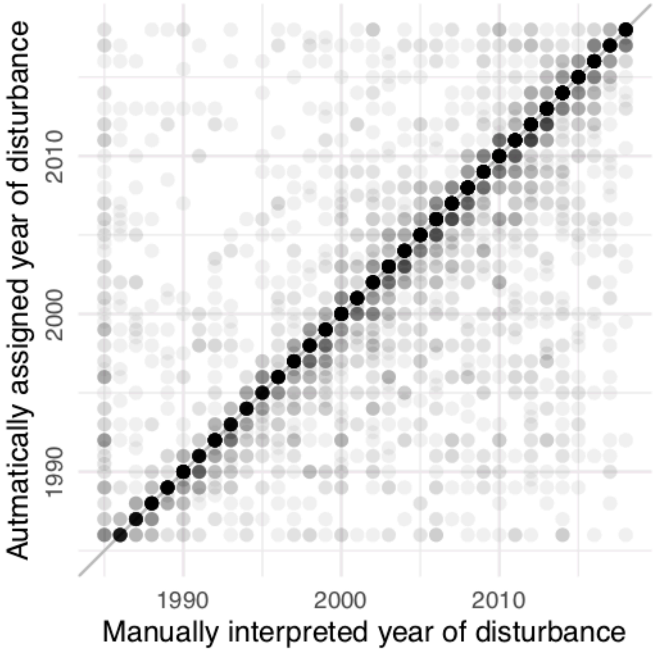


**Figure S5:** Estimated disturbance year versus manually interpreted year of disturbance for 5,000 independent reference pixels. The majority of the pixels is on or close to the 1:1-line, indicating that the correct year of disturbance was assigned. The hue indicates data point density (higher hue = more data points)

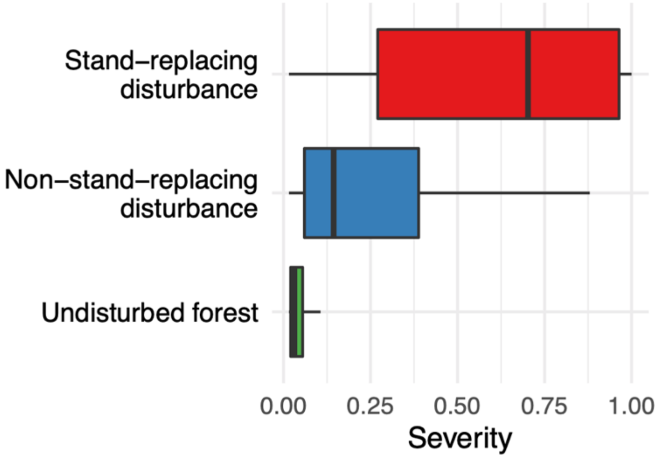


**Figure S6:** Distribution of estimated disturbance severity (i.e., the probability that a pixel/patch has lost its complete canopy during disturbance) among pixels classified as stand-replacing disturbances, non-stand-replacing disturbances and undisturbed forest. The classification labels were derived from reference data and are based on a manual interpretation of Landsat time series and auxiliary use of aerial photos. Stand-replacing disturbances have the highest disturbance severities and are well separated from non-stand-replacing disturbances.

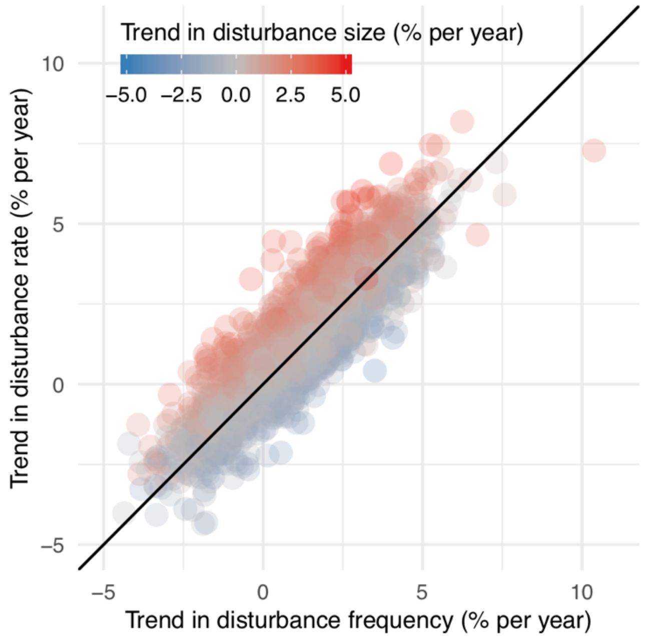


**Figure S7:** Changes in disturbance rates (y-axis; percent of forest area disturbed) in relation to changes in disturbance size (color) and disturbance frequency (x-axis). Trends in disturbance rates are mainly explained by changes in disturbance frequencies (71 %), while changes in disturbance size explained a substantial lower proportion (24 %).

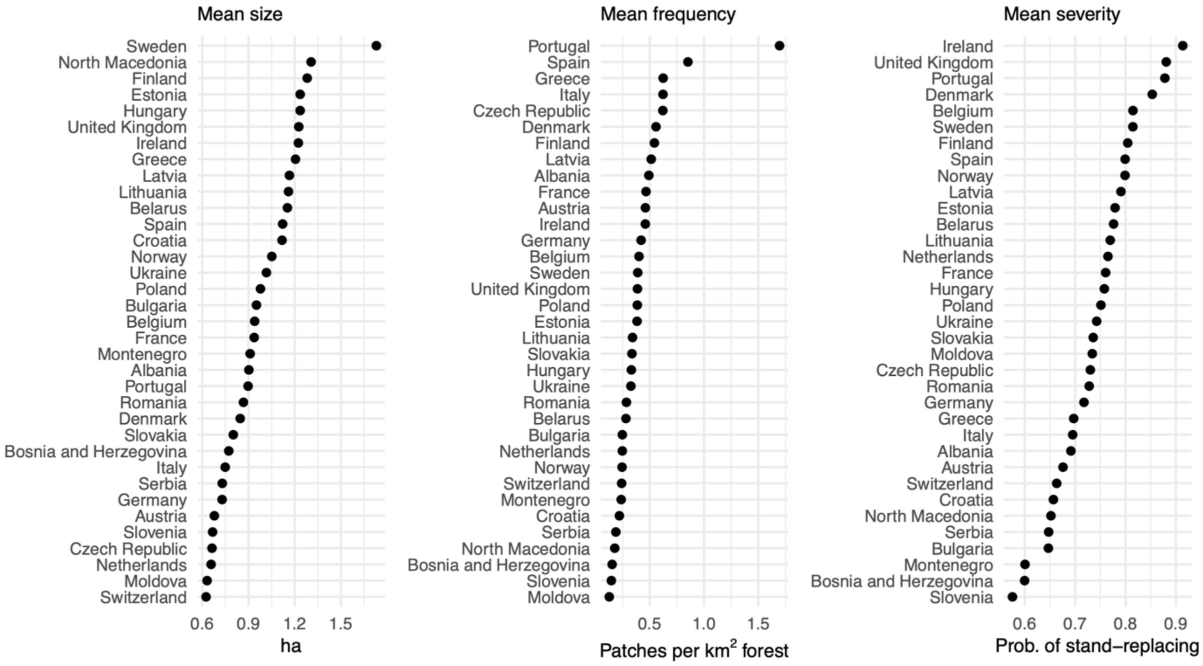


**Figure S8:** Mean disturbance size, frequency and severity summarized for each country of this study. For exact values and other country-wise statistics, please see Table S2.

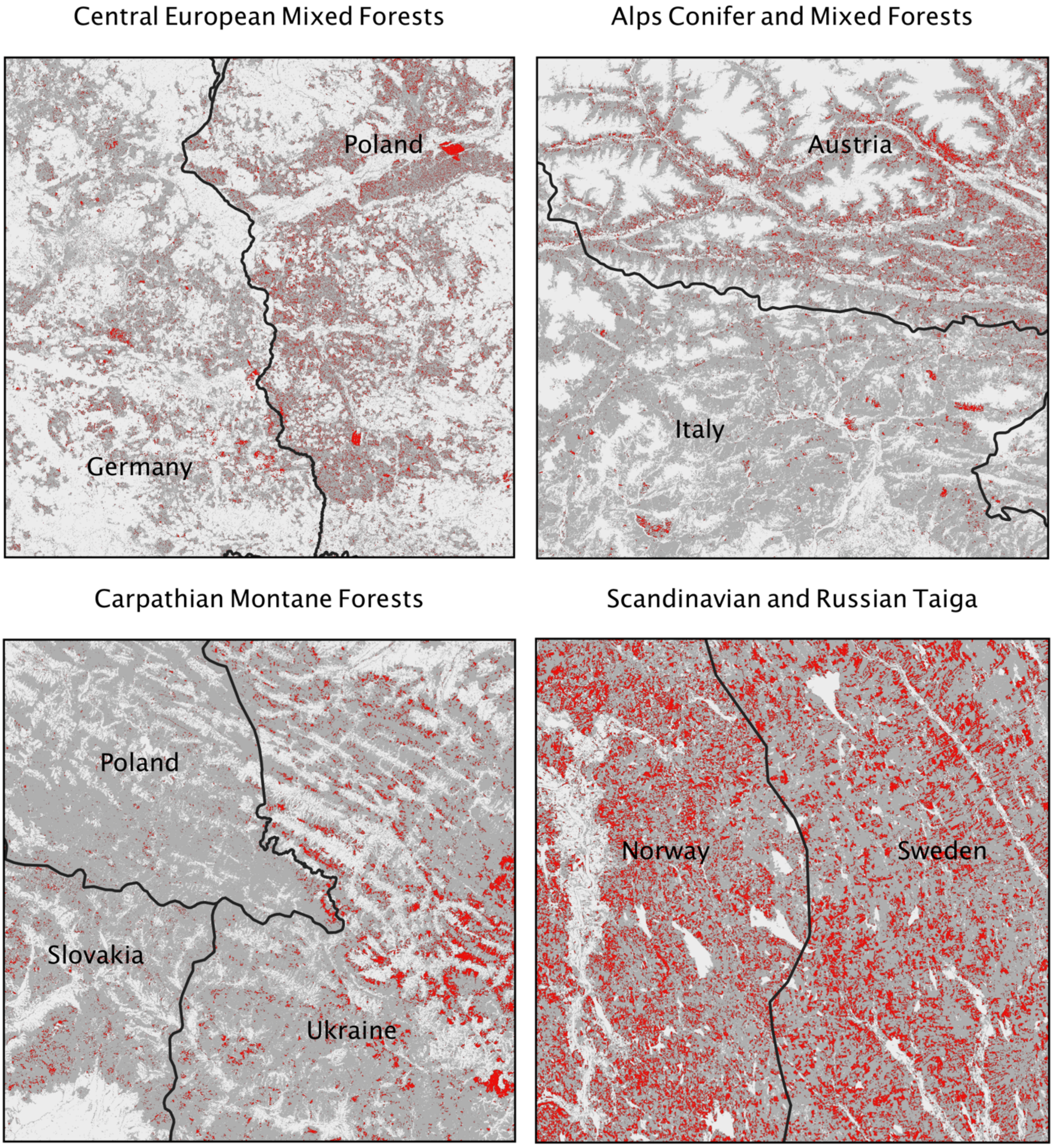


**Figure S9:** Differences in spatial disturbance patterns between countries in similar ecoregions and with similar forest types: (1) Central European Mixed Forests with larger and more frequent disturbances in Poland compared to Germany. (2) Alps Conifer and Mixed Forests with substantially higher disturbance frequencies in Austria compared to Italy. (3) Carpathian Montane Forests, with widely varying disturbances sizes and frequencies between Poland, Slovakia and Ukraine. (4) Scandinavian and Russian Taiga with differences in disturbance size between Norway and Sweden.
