## Supplementary figures and images for "Mapping the forest disturbance regimes of Europe"

### Figure S10

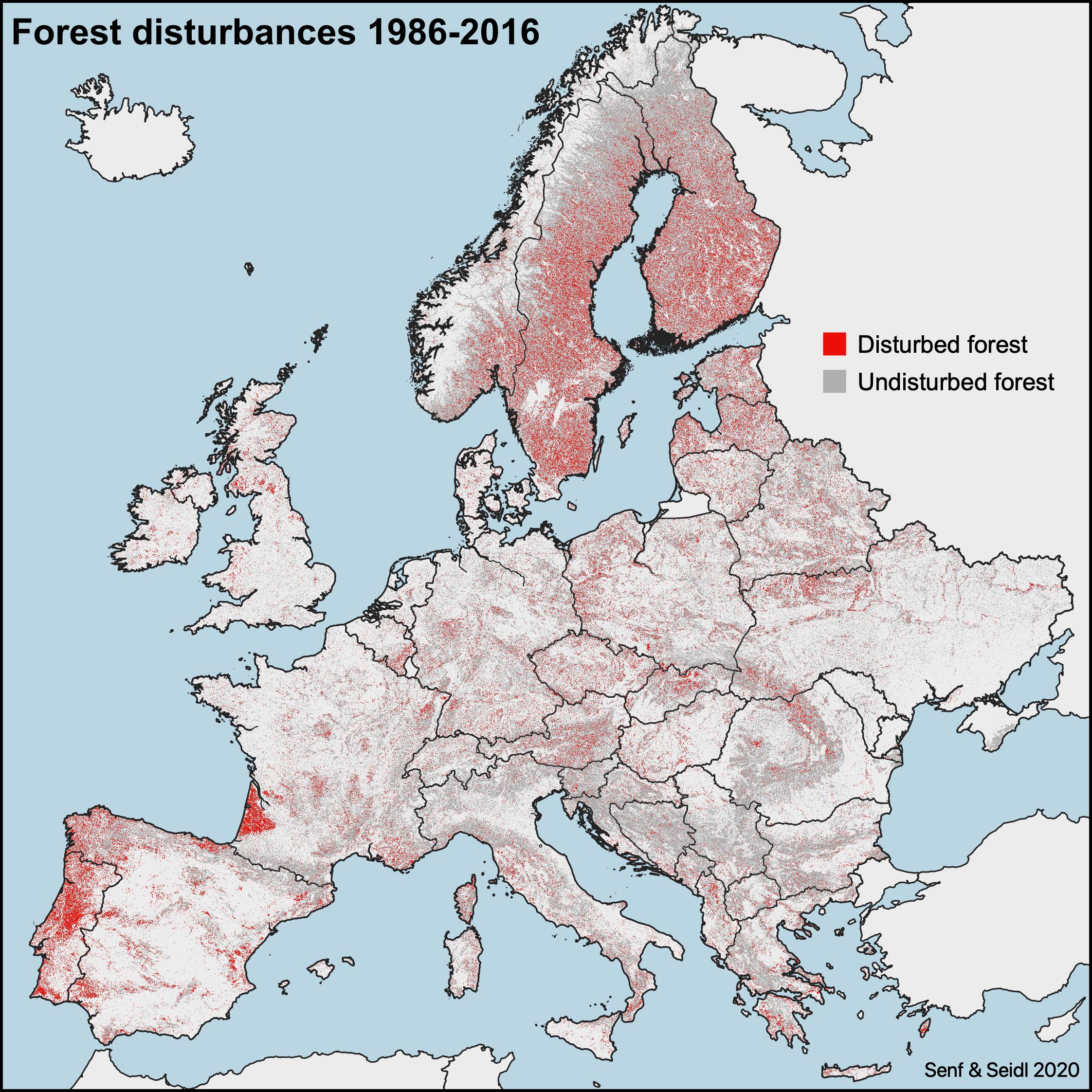
